# Context-dependent reorganization of behavioral strategy during transfer from home-cage to head-fixed performance

**DOI:** 10.64898/2026.01.05.697827

**Authors:** N. Peretz-Rivlin, I. Marsh-Yvgi, Y. Fatal, S. Levin, G. Atlan, A. Citri

## Abstract

Complex behavioral tasks increasingly rely on autonomous training paradigms that enable high-throughput learning in naturalistic settings. Such approaches offer a scalable route for preparing animals for subsequent head-fixed recordings required for high-resolution neural measurements. However, how animals adapt their behavioral strategies across these distinct contexts remains poorly understood. Here, we examine how mice trained autonomously in a group-housed home-cage system reorganize their behavior when transitioned to a head-fixed version of the same delayed-response task. Although overall success rates were comparable across contexts, mice exhibited a striking shift in behavioral policy. Freely behaving home-cage performance was characterized by high engagement and premature responses, whereas head-fixed performance showed a progressive emergence of selective, cue-intensity-dependent responding accompanied by increased omission errors. This selective strategy was strongly associated with improved performance and sustained engagement under head-fixed conditions, reflecting a trade-off between sensitivity to low-intensity cues and suppression of premature actions. Importantly, selectivity did not arise immediately upon head fixation but developed gradually with experience across sessions, indicating learned adaptation rather than a spontaneous response to restraint. Our findings demonstrate that behavioral policy is not solely determined by task rules or sensory demands, but is strongly shaped by task architecture, temporal constraints, and opportunity cost. These results highlight the need to account for context-dependent strategy shifts when interpreting neural activity following transfer from autonomous training to controlled recording environments, and establish a framework for studying adaptive behavior across experimental contexts.

## Introduction

Understanding how neural circuits support complex cognitive functions increasingly relies on complex behavioral paradigms to probe decision-making, impulse control, and selective engagement in rodents (Bari & Robbins, 2013; Carandini & Churchland, 2013). As these tasks grow more sophisticated, incorporating delayed responses, dynamic contingencies, or sensory uncertainty, the demands placed on behavioral training rise accordingly (Aguillon-Rodriguez et al., 2021; Guo et al., 2014). A major bottleneck has therefore emerged: training animals to perform complex tasks remains labor-intensive, slow, and variable across experimenters, limiting scalability and reproducibility (Grieco et al., 2021).

Automated behavioral platforms have begun to address this challenge by enabling high-throughput, standardized training with minimal experimenter intervention (Poddar et al., 2013; Murphy et al., 2020). In particular, home-cage systems allow animals to engage with tasks autonomously, under naturalistic and self-paced conditions that reduce stress and experimenter-induced confounds (Gouveia & Hurst, 2017; Grieco et al., 2021). An emerging but still uncommon strategy is to use autonomous home-cage training as a preparatory stage before transitioning animals to head-fixed recordings. Despite its potential to reduce experimenter workload and improve reproducibility, little is known about how strategies acquired under volitional conditions generalize to the constrained temporal structure of head fixation. This gap is especially relevant for tasks probing cognitive control and behavioral selectivity, where changes in temporal structure and agency may fundamentally reshape strategy and neural encoding.

A useful framework for understanding this question is transfer learning: the application and modification of previously acquired rules in a new context (Pan & Yang, 2010). In animal learning theory, this idea relates closely to learning-set formation, whereby subjects abstract task structure and generalize it across conditions (Harlow, 1949). Rodents are known to exhibit such flexibility, transferring perceptual and decision strategies across sensory modalities, noise levels, and motor contingencies (Busse et al., 2011; Raposo et al., 2012; Siniscalchi et al., 2016). Yet, transfer is rarely cost-free. When task rules are deployed in a new context, animals must reconcile prior strategies with altered sensory feedback, motor demands, and motivational structure, often undergoing a period of behavioral reorganization (Gilad et al., 2018; Bissonette & Powell, 2012). Thus, while several studies have independently examined behavior in freely moving or head-fixed settings, how behavioral strategy reorganizes when animals are transferred from training in a self-paced, freely moving environment to a head-fixed context, while task structure remains unchanged, has not been examined.

This issue is particularly salient for tasks that require the suppression of premature actions and selective responding to relevant cues: core components of cognitive control (Bari & Robbins, 2013b). In such tasks, performance reflects a balance between engagement and restraint: animals must respond reliably to salient signals while withholding responses to ambiguous or noisy inputs (Bogacz et al., 2006; Shenhav et al., 2013). Failures of this balance can manifest as impulsive responses or disengagement, flattening psychometric functions and degrading performance (Berditchevskaia et al., 2016; Aguillon-Rodriguez et al., 2021; Ashwood et al., 2022).

Here, we investigate how behavioral policy reorganizes across experimental contexts by combining autonomous home-cage training with head-fixed task performance. We trained mice in a fully automated, group-housed home-cage system on the ENGAGE task, a delayed-response paradigm that requires response inhibition and auditory cue detection (Peretz-Rivlin et al., 2024; Atlan et al., 2024). Mice acquired the task through volitional, self-paced engagement, generating large and consistent behavioral datasets under naturalistic conditions. The same animals were then transferred to a head-fixed version of the task, enabling direct comparison of behavior across contexts while holding sensory stimuli and task rules constant.

Whether behavioral strategies are preserved or reorganized across contexts remains unknown. Comparing the performance of the same mice while freely behaving vs. while head-fixed, we find that although overall success rates remain comparable, mice undergo a pronounced, experience-dependent shift in behavioral policy. Specifically, head-fixed performance is characterized by the gradual emergence of selective, cue-dependent responding and reduced impulsivity, a strategy that supports sustained engagement and improved performance under constrained trial availability. These findings demonstrate that behavioral policy is strongly shaped by task architecture, even in the absence of changes in task rules, and underscore the importance of accounting for context-dependent strategy shifts when interpreting neural activity following transfer from autonomous training to controlled recording environments.

## Results

### Autonomous learning and volitional participation in the HOMECAGE platform

To examine how behavioral strategies adapt across experimental contexts, we analyzed the behavior of mice trained autonomously in a freely moving, group-housed home-cage system and subsequently transferred to a head-fixed version of the same task. This design allowed us to compare behavior across contexts while holding task rules and sensory stimuli constant. We first characterize volitional task engagement in the home-cage environment, and then examine how behavioral policy reorganizes following transfer to the head-fixed context and across subsequent experience.

To establish a baseline of behavior under fully autonomous, self-paced conditions, we first analyzed task engagement and performance in the home-cage environment. The HOMECAGE accommodates up to five mice per home cage, providing *ad libitum* access to food while restricting water intake to a single behavioral port. This port is equipped with an RFID reader to identify individual animals during task engagement (Figure 1A) (Peretz-Rivlin et al., 2024). We utilized the HOMECAGE system to train mice on the ENGAGE task, a delayed-response paradigm that combines response inhibition with auditory cue detection. Trials were initiated upon RFID detection of the mouse within the operant tube. A trial onset *WAIT*-signal (100 ms broadband noise) indicated the start of a variable delay period (0.75–2.5 sec), during which mice were required to withhold responding. Following this delay, an auditory *GO*-cue (6 kHz, 5 pulses, 100 ms duration) signaled the start of a 1.5 s response window. A response during this window triggered the delivery of a reward (4 µl water). To modulate task difficulty, the *GO*-cue was presented at four logarithmically scaled intensities. Additionally, 20% of trials included a simultaneous visual aid (LED at the lick port) to facilitate cue detection, while 50% of trials featured the *GO*-cue embedded within a background noise tone cloud (Figure 1B). Mice had continuous, 24-hour access to the operant interface, allowing for fully volitional scheduling. We observed that task engagement followed natural circadian rhythms, with activity predominantly concentrated during the dark cycle (Figures 1C, D). Furthermore, interactions with the operant tube occurred in a clustered manner, characterized by peak of short inter-trial intervals (<1 min) that indicate sustained behavioral bouts (Figure 1E). These patterns indicate that task engagement in the home-cage environment is both volitional and self-paced, allowing mice to initiate trials primarily during periods of high motivation. This structure minimizes opportunity cost for impulsive actions, a feature that becomes central when comparing behavior across contexts.

**Figure 1.**
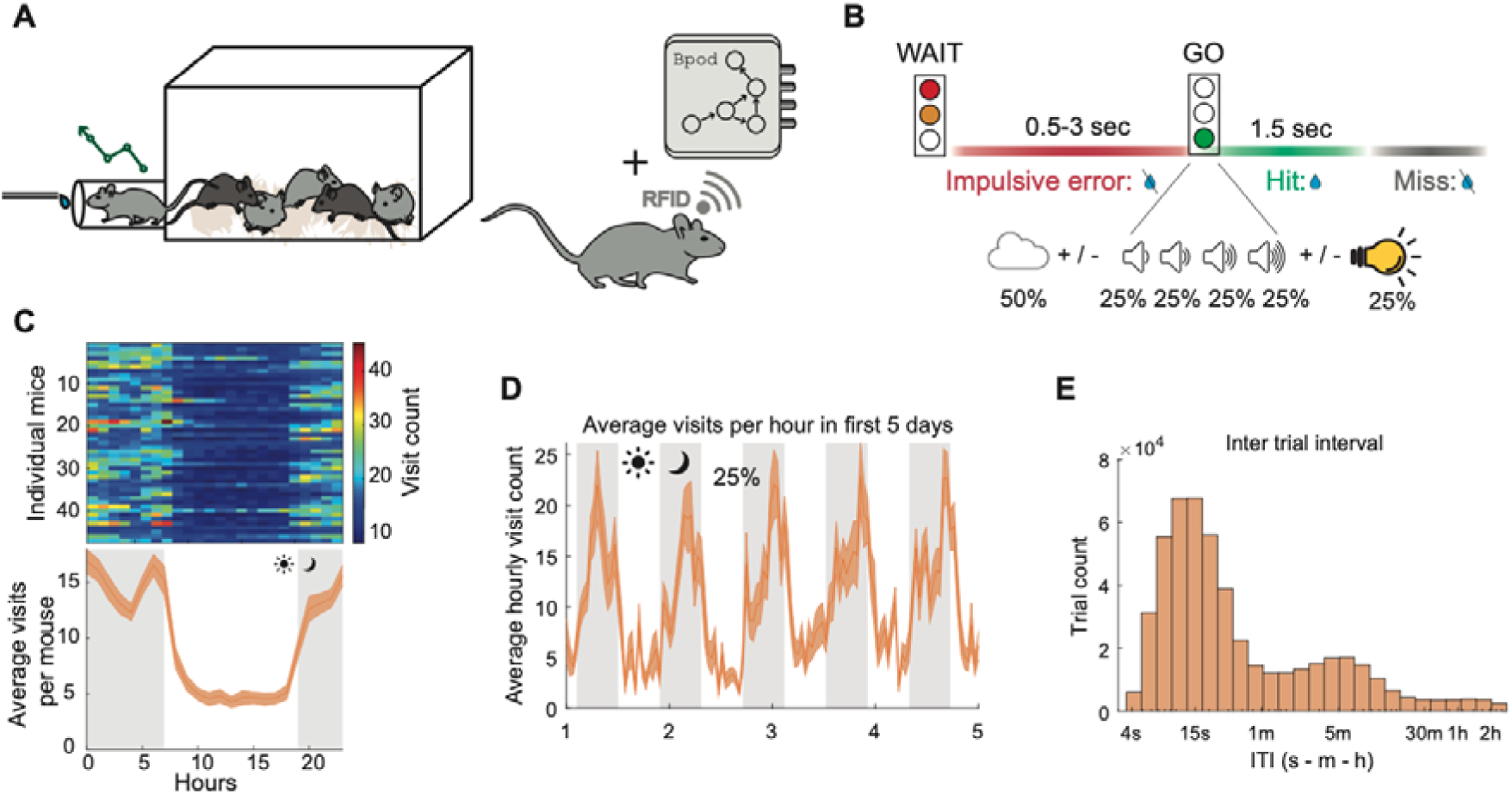
Volitional engagement of mice in the HOMECAGE system enables autonomous learning of the ENGAGE task. **(A)** Schematic illustration of the HOMECAGE system and its main components. **(B)** Structure of the ENGAGE task. Each trial began with a *WAIT* signal instructing mice to withhold responding for a variable delay period, after which the *GO* cue was presented. The *GO* cue consisted of one of four equally probable sound intensities and was presented atop a background tone cloud in 50% of trials. In 25% of trials, the *GO* cue was accompanied by a visual aid (an LED within the poke port). **(C)** *Top:* Heatmap showing the distribution of operant port visits across hours of the day, averaged across training days; each row corresponds to a single mouse. *Bottom:* Mean hourly visit rate across mice (solid line), with shaded area representing SEM (n = 47 mice). **(D)** Average hourly visit rate during the first five days of access to the HOMECAGE system; line represents the across-mouse mean (n = 47 mice). **(E)** Distribution of inter-trial intervals (ITIs) across all mice (n = 47), displayed on a semi-logarithmic x-axis.

### Training procedure

Mice acquired the task through a structured, multi-stage shaping curriculum designed to incrementally increase difficulty. Initially, training focused on associating the operant port with water availability; in this phase, every poke was rewarded, and water delivery was paired with both the auditory *GO*-cue and a visual aid to facilitate cue-outcome association. Subsequently, a response inhibition component was introduced, requiring mice to wait for the *GO*-cue during a short delay (0.5–2.0 s). In the third stage, the delay period was extended (randomized 0.75–2.5 s), and the visual aid was gradually withdrawn to enforce reliance on auditory cues. Finally, task complexity was increased by introducing background noise, initially scaffolded by the visual aid before the aid was removed entirely, and by varying the *GO*-cue across four sound intensities (Figure 2A).

**Figure 2.**
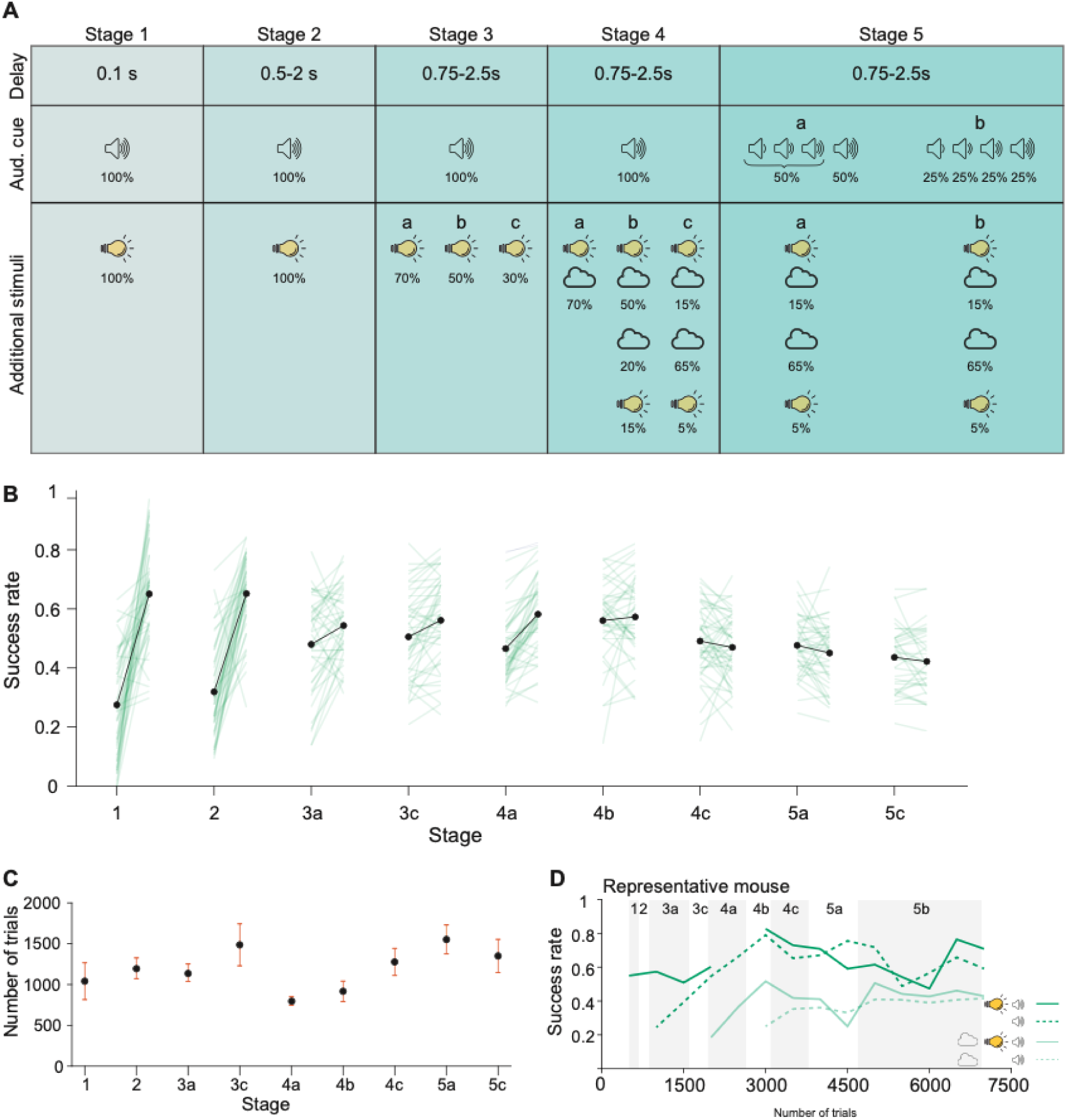
Training procedure and acquisition of the ENGAGE task. **(A)** Stepwise training protocol for acquiring the ENGAGE task. Task components were introduced progressively: the delay period was extended across stages, the proportion of *GO* cues without a visual aid was increased, the background tone cloud was added in stage 4, and varying sound intensities were introduced in stage 5. Transitions between stages were triggered based on consistency in performance. **(B)** Success rates across the training phase. For each stage, performance in the first 15% of trials is compared to the last 15%. Green lines represent individual mice (n = 47); the black line indicates the across-mouse mean. **(C)** Number of trials completed per training stage. Dots represent the mean across mice (n = 47); error bars indicate SEM. **(D)** Representative task acquisition for an individual mouse across trial types over the course of training. Dark green traces represent trials without a background tone cloud; light green traces represent trials with a tone cloud. Solid lines denote trials with a visual aid, and dashed lines denote trials without a visual aid.

Progression through the training curriculum was managed programmatically, tailored to the performance of individual animals. The RFID-based identification system ensured that each mouse received the specific task protocol corresponding to its current training stage, regardless of the status of its cage-mates. In early training phases, performance was characterized by a rapid increase in success rates from stage onset to completion (Figure 2B). As task complexity increased, mice reached stable but imperfect performance, ensuring sufficient error trials for subsequent behavioral and physiological analyses. Training duration varied substantially across individuals, highlighting natural variability in learning trajectories (Figure 2C). A representative learning trajectory (Figure 2D) illustrates this trend: a sharp increase in success rates during initial acquisition followed by stable performance plateaus as task difficulty escalated.

### Context-dependent emergence of behavioral selectivity in the ENGAGE task

Following autonomous task acquisition in the home-cage environment, mice were transferred to a head-fixed setup to enable neurophysiological recording (Figure 3A). We implemented a rapid transfer protocol that reliably yielded task engagement within three sessions. After removal from the home cage, mice underwent a brief period of water restriction before an initial head-fixed session focused on reacclimating the motor response. During this first session, water reward was replaced with sucrose solution, and either Pavlovian auto-rewards (reward co-delivered with the *GO-*cue) or a simplified task version (high-intensity cues without background noise) was used to promote engagement. By the second session, mice were performing the full task, and stable behavior was typically achieved within one to two habituation sessions prior to recording.

**Figure 3.**
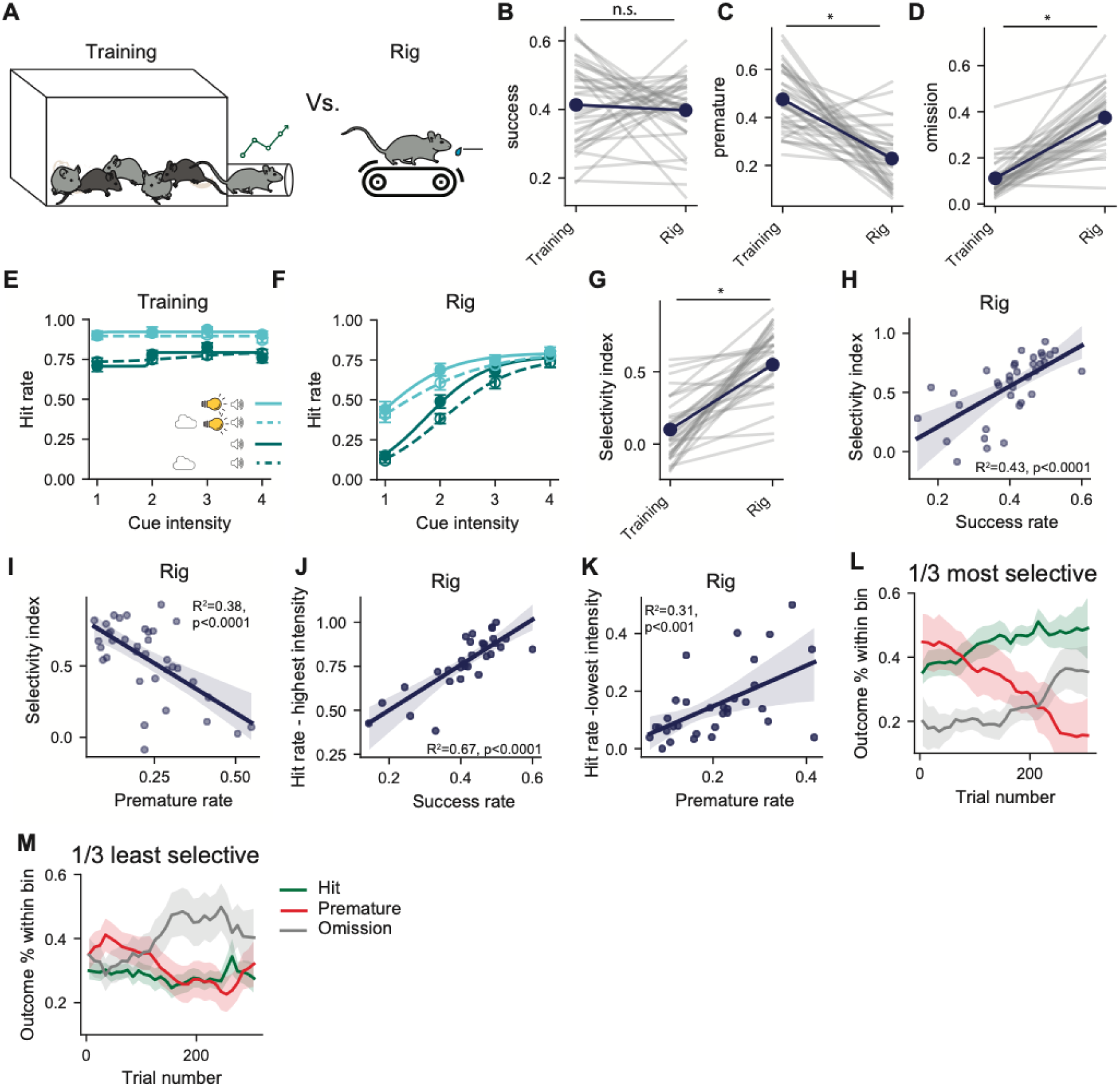
Behavioral selectivity increases as mice transition from autonomous training in the automated cage to head-fixed performance. **(A)** Schematic illustration of the behavioral contexts in which mice learned and performed the ENGAGE task: the HOMECAGE automated system and the head-fixed rig. **(B)** Success rates during the final stage of home-cage training compared with performance in the head-fixed rig. Gray lines represent individual mice (n = 36); navy points and line represent the across-mouse mean. **(C)** Premature response rates in the two contexts. Gray lines indicate individual mice (n = 36); navy points and line indicate the mean (t=7.153, p<0.0001 paired t-test). **(D)** Omission error rates (t=-10.780, p<0.0001 paired t-test; n = 36 mice). **(E)** Psychometric performance curves in the home-cage system, showing hit rate as a function of cue intensity and cue type. Cyan: trials with a visual aid; dark green: trials without a visual aid. Solid lines denote trials without a background tone cloud; dashed lines denote trials with a tone cloud (n = 36 mice). **(F)** Corresponding psychometric curves for the head-fixed setup. **(G)** Selectivity Index (SI; difference between hit rate for the highest-intensity cue and the lowest-intensity cue) in the two behavioral setups (t=-9.036, p<0.0001 paired t-test; n = 36 mice). **(H)** Relationship between SI and success rate during head-fixed testing sessions (simple linear regression; R² = 0.43, p < 0.0001, n = 36 mice). **(I)** Relationship between SI and premature response rate during head-fixed testing sessions (R² = 0.38, p < 0.0001, n = 36 mice). **(J)** Hit rate for the highest-intensity cue (with visual aid) plotted against success rate in head-fixed sessions (simple linear regression; R² = 0.67, p < 0.0001, n = 36 mice). **(K)** Hit rate for the lowest-intensity cue plotted against premature response rate during head-fixed testing (simple linear regression; R² = 0.31, p < 0.001; n = 36 mice). **(L)** Proportion of trial outcomes (correct = green, premature = red, omission = gray) across behavioral sessions for the top one-third most selective mice. Lines represent across-session means; shaded areas represent SEM (n = 12 mice). **(M)** Same analysis as in (L) for the one-third least selective mice (n = 12 mice).

Importantly, while the sensory stimuli and task structure were retained across contexts, the environments differed fundamentally in their temporal and motivational architecture. In the home cage, trial initiation was fully volitional and self-paced, with no imposed inter-trial interval. In contrast, head-fixed sessions consisted of discrete, time-limited blocks of trials initiated externally at fixed inter-trial intervals (18-20 sec), under conditions of physical restraint.

Despite these differences, overall task performance was comparable across contexts. Success rates during the final stage of home-cage training closely matched those observed during head-fixed sessions (Figure 3B), indicating that mice retained task knowledge following transfer. However, the structure of errors differed markedly between environments. In the home-cage context, error trials were dominated by premature responses during the delay period following the *WAIT* signal, reflecting a high propensity for impulsive responses (Figure 3C). In contrast, head-fixed sessions exhibited substantially fewer premature responses but a pronounced increase in omission errors, in which mice failed to respond during the response window (Figure 3D). This shift suggests that a trade-off between impulsivity and disengagement emerges under constrained task conditions.

These differences were further reflected in psychometric performance. In the home-cage environment, response rates were uniformly high across cue intensities and task conditions, resulting in relatively flat psychometric functions (Figure 3E). Hit rates showed minimal modulation by *GO*-cue intensity, background noise, or the presence of a visual aid, indicating low selectivity across stimulus conditions. In contrast, head-fixed performance exhibited sigmoidal psychometric functions: response probability scaled with cue intensity, was enhanced by the visual aid, and was attenuated by background noise (Figure 3F). Because auditory stimuli and cue intensities were identical or amplified in the head-fixed setup, the emergence of selectivity is unlikley to reflect improved sensory sensitivity and instead indicates a reorganization of response strategy.

To quantify this shift, we computed a Selectivity Index (SI; defined as the difference in hit rates between the highest- and lowest-intensity *GO* cues). SI was significantly higher in the head-fixed context than during home-cage training (Figure 3G), capturing the emergence of selective responding under constrained conditions. This increase in selectivity proved behaviorally relevant for task performance. Across animals, the SI was strongly and positively correlated with overall success rate during head-fixed sessions (Figure 3H), indicating that more selective mice achieved higher performance under constrained task conditions. Decomposing performance by error type revealed how selectivity relates to distinct classes of errors. Higher selectivity was associated with significantly lower rates of premature responses during head-fixed sessions (Figure 3I), indicating that selective mice were more effective at withholding responses during the delay period. This reduction in premature responding points to improved response control rather than a generalized decrease in engagement.

Further decomposing hit rates by *GO*-cue intensity revealed a clear dissociation between responses to strong and weak cues. Hit rates for the highest-intensity cues were tightly correlated with overall success rate (Figure 3J), indicating that high-performing, selective mice reliably detected and responded to salient task-relevant signals. In contrast, higher hit rates for the lowest-intensity cues were associated with increased premature responding (Figure 3K), suggesting that attempts to respond to near-threshold stimuli were coupled to impulsive behavior. Together, these results indicate that behavioral selectivity reflects a strategic trade-off: prioritizing responses to robust sensory evidence while suppressing responses to ambiguous cues supports higher task efficiency under constrained conditions.

Selective and non-selective response strategies diverged markedly over the course of a behavioral session. To characterize this divergence, mice were stratified into tertiles based on their SI, and the temporal distribution of trial outcomes was examined across head-fixed sessions. Striking differences emerged in the session dynamics of highly selective versus weakly selective animals (Figures 3L, M). Both groups began sessions with elevated rates of premature responses, consistent with high initial motivation following water restriction. In highly selective mice, premature responding declined progressively over the session, accompanied by a corresponding increase in correct responses that consistently exceeded omission errors (Figure 3L). This pattern reflects a transition into a stable and efficient performance state characterized by sustained engagement and effective response control. In contrast, mice with low selectivity failed to enter this stable regime. Although premature responding also declined over time in this group, it was accompanied by a marked increase in omission errors rather than an increase in correct responses (Figure 3M), indicating rapid disengagement from the task.

These divergent trajectories suggest that selective responding supports sustained engagement under constrained task conditions, whereas non-selective strategies, characterized by attempts to respond to weak or ambiguous cues, accelerate disengagement and limit performance across the session.

### Experience-dependent development of selective behavior in the head-fixed context

The analyses above indicate that selective responding supports efficient performance and sustained engagement during head-fixed sessions. Notably, selective responding did not emerge immediately upon transfer to the head-fixed context but instead developed with experience. To quantify this process, we examined behavioral performance across the initial head-fixed sessions, separating an early habituation phase (the first two sessions) from subsequent test sessions in which behavior stabilized (Figure 4A). Transition from the home-cage environment to the recording rig induced a transient reduction in success rate during habituation, which recovered during later sessions (Figure 4B). In contrast, premature response rates declined rapidly following transfer and continued to decrease across test sessions (Figure 4C), indicating that impulsive responding is rapidly suppressed following transition to head-restraint and associated shifts in task structure.

**Figure 4.**
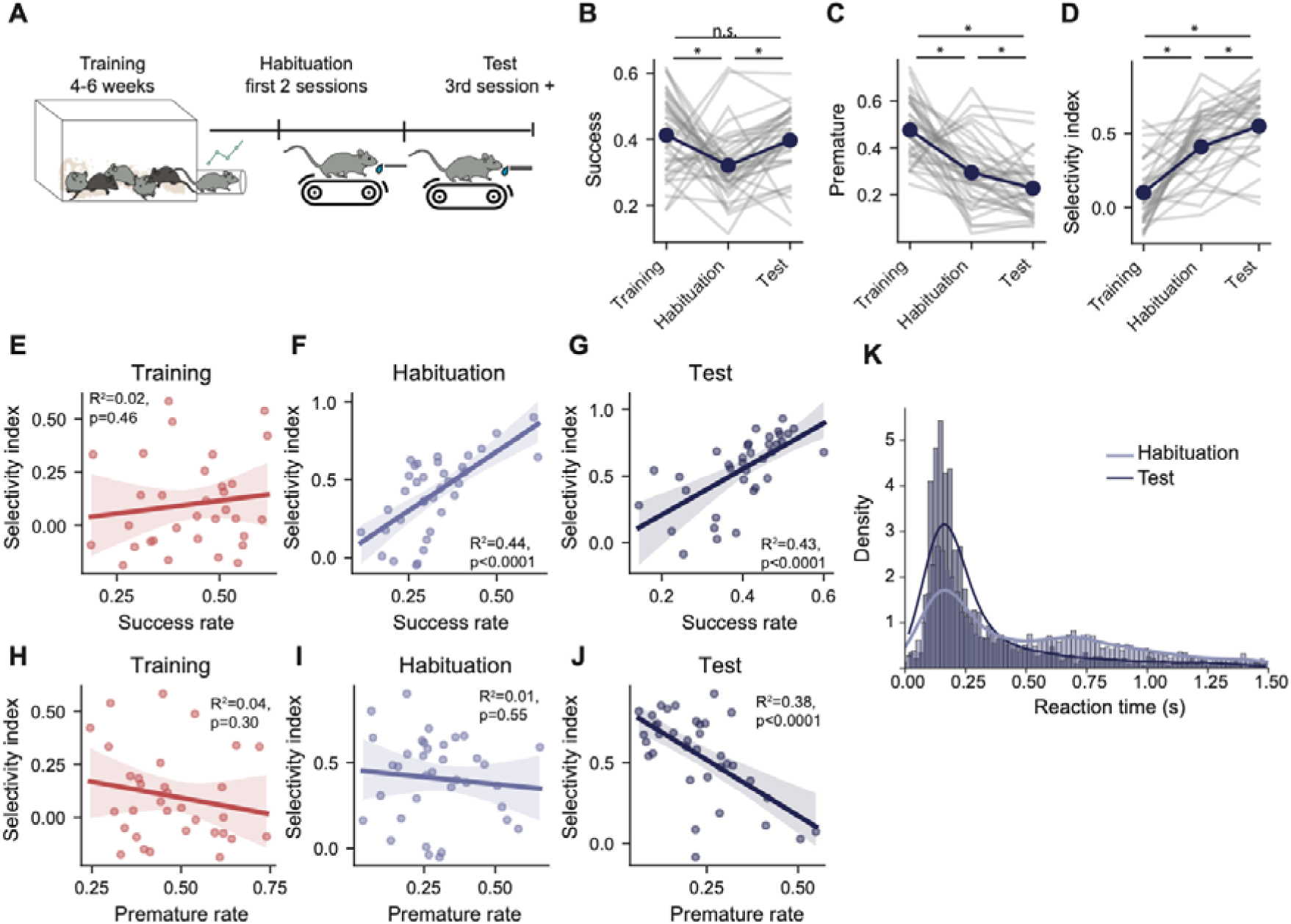
Gradual emergence of selective behavior across training, habituation, and test sessions in the head-fixed context. **(A)** Schematic overview of the transition from autonomous training in the HOMECAGE system to head-fixed behavioral performance. Following home-cage training, mice underwent two habituation sessions in the head-fixed rig; test sessions began on the third head-fixed session. **(B)** Success rates across training, habituation, and test phases. Gray lines represent individual mice (n = 36); the navy line indicates the across-mouse mean. Success rates during habituation were significantly lower than during home-cage training (p = 0.018) and test sessions (p < 0.001), whereas training and test did not differ (p = 0.588). Statistics reflect Bonferroni-corrected post hoc t-tests following a mixed-effects linear model. **(C)** Premature response rates across phases. Premature errors decreased from training to habituation (p < 0.0001) and further decreased from habituation to test sessions (p = 0.013). **(D)** SI across phases. SI increased from training to habituation (p < 0.0001) and increased further from habituation to test sessions (p < 0.0001). **(E)** Relationship between SI and success rate during training (simple linear regression; R² = 0.02, p = 0.46; n = 36 mice). **(F)** Relationship between SI and success rate during habituation (simple linear regression; R² = 0.44, p < 0.0001; n = 36 mice). **(G)** Relationship between SI and success rate during test sessions R² = 0.43, p < 0.0001; n = 36 mice). **(H)** Relationship between SI and premature response rate during training (simple linear regression; R² = 0.04, p = 0.30; n = 36 mice). **(I)** Relationship between SI and premature response rate during habituation (simple linear regression; R² = 0.01, p = 0.55; n = 36 mice). **(J)** Relationship between SI and premature response rate during test sessions (simple linear regression; R² = 0.38, p < 0.0001(; n = 36 mice). **(K)** Distribution of reaction times during habituation and test phases.

The SI, however, followed a distinct trajectory. SI increased during the habituation phase and also continued to rise significantly during subsequent test sessions (Figure 4D), indicating that selective, cue-dependent responding is not an immediate consequence of head fixation but instead develops gradually with experience. Consistent with this interpretation, the relationship between SI and performance evolved across phases. During home-cage training, SI showed no meaningful association with either success rate or premature responding (Figures 4E, H). A positive correlation between SI and success rate (but not premature errors) emerged during habituation (Figure 4F, I), whereas a strong negative correlation between SI and premature responses was only evident during test sessions (Figures 4G, J).

Reaction time analysis further supported the emergence of a more efficient behavioral strategy. As mice progressed from habituation to test sessions, reaction time distributions became sharper, with a reduction in long-latency responses (Figure 4K), consistent with improved temporal precision and response control. Together, these findings demonstrate that while head fixation immediately alters aspects of behavior such as impulsivity, the selective strategy that supports efficient performance is acquired gradually through experience in the constrained task environment. Thus, suppression of impulsive responding and emergence of selective cue dependence follow dissociable time courses, indicating that selectivity reflects learning rather than a direct consequence of physical restraint.

Collectively, the reported observations demonstrate that behavioral policy is not fixed by task rules alone but is adaptively reorganized through experience when learned behavior is deployed under altered temporal and motivational constraints.

## Discussion

The results presented here show that transferring identical task rules from a freely moving to a head-fixed context induces a learned reorganization of behavioral policy. Although mice retained task knowledge and achieved comparable overall success rates across contexts, they adapted how that knowledge was deployed. Freely moving home-cage performance was characterized by high engagement and frequent premature responses, whereas head-fixed performance was marked by the gradual emergence of selective, cue-dependent responding that supported efficient performance and sustained engagement. Unlike classical learning-set formation, which involves abstraction of task rules, the transfer observed here reflects reorganization of how known rules are deployed under altered temporal and motivational constraints. These findings demonstrate that behavioral policy is not fixed by task rules alone, but is flexibly shaped by the temporal and motivational structure of the environment.

In the home-cage environment, trial initiation is entirely volitional: mice engage with the task when motivated, and nearly every initiated trial includes a response. Under these conditions, premature actions carry little cost, as an impulsive response simply resets the trial and allows an immediate retry. In contrast, the head-fixed environment imposes externally paced trials with long, fixed inter-trial intervals and a limited session duration. In this setting, premature responses incur a substantial opportunity cost, by forfeiting reward and consuming a finite trial opportunity, thereby biasing animals toward more conservative strategies that suppress premature responding.

Temporal constraints further shape this adaptation. In the home cage, mice can distribute effort flexibly across the circadian cycle, engaging with the task when motivational and energetic states are favorable. By contrast, head-fixed sessions compress hundreds of trials into a tightly bounded time window, making sustained engagement energetically demanding. Under these conditions, ignoring ambiguous or low-certainty stimuli and responding preferentially to robust cues may support more efficient allocation of effort, a pattern reflected in the evolving distribution of trial outcomes across sessions. Together, these observations highlight how task architecture, independent of sensory stimuli or explicit rules, can exert a strong influence on behavioral strategy.

The emergence of selective behavior raises important questions about underlying mechanisms. One possibility is that selectivity reflects changes in response threshold or engagement state: animals adopting lower thresholds respond to weak cues but incur more premature errors, whereas higher thresholds promote selectivity at the cost of increased omissions. A complementary possibility is that sensory representations themselves differ across contexts, with more reliable encoding of high-intensity cues facilitating rapid, automatic responses under head fixation. Disentangling these possibilities will require future experiments that combine behavioral analysis with neural recordings and causal manipulations to determine whether selectivity arises primarily from shifts in responsivity, sensory processing, or downstream decision circuitry.

Beyond the specific task examined here, our findings have broader implications for experimental design. Automated home-cage training systems provide powerful tools for scalable, low-confound behavioral acquisition, but transferring animals to head-fixed recording environments introduces fundamental changes in temporal structure and opportunity cost that can reshape behavioral policy. Accounting for these changes is essential for interpreting neural activity measured after transfer. At the same time, this framework presents a unique opportunity: by holding task rules constant while altering environmental constraints, it enables systematic investigation of how learned behavior is reconfigured across contexts. Such analyses expose meaningful individual differences in adaptive strategy use and provide a window into the neural mechanisms that support flexibility, strategy selection, and context-sensitive control.

In conclusion, this work establishes a reproducible behavioral framework that links autonomous, naturalistic learning with controlled head-fixed task performance, revealing how behavioral policy emerges from the interaction between learned task structure and environmental constraints. By demonstrating that selective, cue-dependent responding is acquired gradually through experience in the head-fixed context, these findings set the stage for future studies that integrate large-scale recordings and causal interventions to uncover the neural computations that support adaptation and strategic control across behavioral environments.

## Methods

### Animals

In this work we combine data from several cohorts that participated in several experiments (see Supplementary Table 1 for details of mice analyzed under both freely-behaving and head-fixed conditions). All key effects were replicated independently within each cohort (data not shown). A total number of 47 mice underwent automated training, of which we present 36 mice that also transitioned to the behavioral setting. 11 mice were excluded from the analysis of performance on the rig as they were subject to manipulations that directly altered task engagement or response policy and could therefore not be meaningfully merged with unmanipulated animals. All mice described in this study were male C57BL/6JOLAHSD obtained from Harlan Laboratories, Jerusalem, Israel (RRID:IMSR_JAX:000664). Mice, ages 8-12 weeks old, 22-26 grams at the beginning of the experiment were housed in groups of littermates and kept in a SPF (specific pathogen free) animal facility under standard environmental conditions: temperature (20–22 °C), humidity (55 ± 10%), and 12–12 h light/dark cycle, with ad libitum access to water and food. All experimental procedures, handling, surgeries, and care of laboratory animals used in this study were approved by the Hebrew University Institutional Animal Care and Use Committee (NS-19-15584-3; NS-19-15788-3, NS-20-16301-4).

### Inclusion/exclusion criteria, blinding & power justification

No animals or sessions were excluded based on performance. Experimenters were aware of mouse identity; however, behavioral metrics were defined a priori and computed using automated analysis pipelines applied uniformly across animals. Given the integrative nature of the study, no formal a priori power calculation was performed; observed effect sizes and confidence intervals are reported to support interpretation.

### System Control and Software Architecture

Experimental control and data acquisition were managed by a Bpod State Machine (Sanworks, Model 102; RRID:SCR_015943) interfaced with the Bpod Gen2 software environment in MATLAB (RRID:SCR_001622). To enable individualized training within a group-housed setting, we used ‘Bpod’s “softcode” module to integrate RFID identification into the state machine logic. Specifically, the final state of each trial triggers a softcode byte that initiates a polling loop, repeatedly querying the RFID reader until a valid tag is detected. Upon identification of the subject, the software retrieves the mouse-specific performance history and loads the appropriate task parameters (e.g., current training stage, cue difficulty) for the subsequent trial. This architecture ensures that trial initiation is strictly gated by identification, allowing for autonomous, individualized progression through the training curriculum.

### ENGAGE behavioral task

Training in preparation for fiber photometry or Neuropixels recording in head-fixed mice was performed in automated behavioral cages (see Automated behavioral training below). Subsequently, trained mice were habituated to head-fixation and performance of the ENGAGE task while head-fixed but free to run on a linear treadmill (Janelia 2017-049 Low-Friction Rodent-Driven Belt Treadmill). Each trial in the task is initiated with a brief auditory broad-band noise stimulus to which the mice must refrain from responding (‘*WAIT’*), followed by an auditory ‘*GO’* cue (a set of five 6 kHz pure tone pips). A delay period of random duration (0.5-3 seconds) separates the ‘*WAIT’* from the ‘*GO’* cues. Licks during this delay period (i.e. between the ‘*WAIT’* and the ‘*GO’* cues) result in the termination of the trial and label the trial result as ‘premature response. Following the ‘*GO’* cue, a 1.5-second-long response window is open for a correct lick (‘Hit’). Absence of licking within the defined response window labels the trial as a ‘omission’ and is not rewarded. ‘Hit’ trials are rewarded with a single drop (4 µl) of sucrose water. Trial difficulty was determined by a combination of several factors: four equally probable intensities of the ‘*GO’* cue tone, inclusion of a tone-cloud ‘mask’ (4 seconds of continuous chords assembled from logarithmically spaced pure tones in the frequency range of 1-10 kHz, excluding the *GO* cue frequency, in 50% of trials;), and, in 20% of trials, a visual stimulus (‘visual aid’) presented concurrently with the ‘*GO’* cue. In a subset of the mice used as a control group in au optogenetic experiment there was no visual aid (see Supplementary table 1 for details). Trials were initiated automatically every 18-20 seconds [20 seconds for cohort A (n = 23 mice), published in Atlan et al., 2024; 18 seconds for cohort B (n = 13 mice)]. In cohort A (n = 23) the delay on the rig was randomized between 0.75 to 3 sec, while in cohort B (n = 13 mice) the delay was randomized between 0.75 to 2.5 sec. Analyses pooling head-fixed sessions were replicated within each cohort yielding comparable results; ITI and delay time were therefore not treated as a grouping variable. Behavioral sessions contained blocks of trials containing 15 occurrences (in some cases shortened to 8 or 10) of each possible combination of parameters, in random order. These blocks were repeated 2–4 times (as long as mice maintained participation) for a total of up to 1,000 trials per mouse per day (a typical daily session totaled 240-480 trials).

### Behavioral Shaping and Curriculum. Mice were trained on the ENGAGE task using an automated, multi-stage shaping curriculum

Training began with a “lick adaptation” phase, where mice learned to associate auditory-visual cues with water availability. Following acquisition of this association, mice progressed through five stages of increasing difficulty, with transitions triggered based on performance criteria (typically stable behavior at 50–70% accuracy).

**Stage 1**: Introduction of the basic task structure with a short random delay. (Average duration: 2.6 days).

**Stage 2**: The random delay period was extended to 0.5–2.0 s. (Average duration: 2.5 days).

**Stage 3**: Introduction of the full delay range (0.5–2.5 s) and the gradual withdrawal of the visual aid. This fading occurred in three sub-stages, reducing the proportion of visually aided trials to 30%, 50%, and finally 70% auditory-only trials. (Average duration: 4.4 days).

**Stage 4**: Introduction of a background masking noise (Tone Cloud; 1–10 kHz chords, 67.5 dB SPL).

Stage 5: Introduction of additional three distinct target cue attenuations to probe perceptual sensitivity.

Throughout all stages, premature or late licks were unrewarded and required the mouse to exit the port, terminating the RFID read, and re-enter to initiate a new trial. Total training time to expert performance averaged 29.6 days.

### Stereotactic surgery and viral injections

Induction and maintenance of anesthesia during surgery were achieved using SomnoSuite LowFlow Anesthesia System (Kent Scientific Corporation). Following induction of anesthesia, animals were quickly secured to the stereotaxic apparatus (David KOPF instruments). Anesthesia depth was validated by toe-pinching and manual heart-rate monitoring. Isoflurane levels were adjusted (0.8–1.5%) to maintain a heart rate of ∼60 bpm. The skin was cleaned with Betadine (Dr. Fischer Medical); Lidocaine (Rafa Laboratories) was applied to minimize pain; and Viscotears gel (Bausch & Lomb) was applied to protect the eyes. An incision was made to expose the skull, which was immediately cleaned with hydrogen peroxide, and a small hole was drilled using a fine drill burr (model 78001RWD Life Science). Using a microsyringe (33GA; Hamilton syringe) connected to an UltraMicroPump (World Precision Instruments), a virus was subsequently injected at a flow rate of 50–100 nl/min, after which the microsyringe was left in the tissue for 5–10 min after the termination of the injection before being slowly retracted. For photometry or optogenetic experiments, a fiberoptic ferrule (400 μm, 0.37–0.48 NA, Doric Lenses) was slowly lowered into the brain. A custom-made metal head bar was glued to the skull, the incision was closed using Vetbond bioadhesive (3M), and the skull was covered in dental cement and let dry. An RFID (radio-frequency identification) chip (ID-20LA, ID Innovations) used for tracking during behavioral training was implanted subcutaneously. Mice were then disconnected from the anesthesia and were administered a subcutaneous saline injection for hydration and an IP injection of the analgesic Rimadyl (Norbrook) as they recovered under gentle heating. Coordinates for the claustrum were based on the Paxinos and Franklin mouse brain atlas. Unless noted otherwise, viruses were prepared at the vector core facility of the Edmond and Lily Safra Center for Brain Sciences at the Hebrew University. See Supplementary Table 1 for injection sites and viruses used.

### Data and code availability

Behavioral data, task control code and parameters, as well as analysis code will be deposited in a public repository upon acceptance for review (GitHub/Zenodo; DOI to be provided).

---

**Supplementary table 1:**
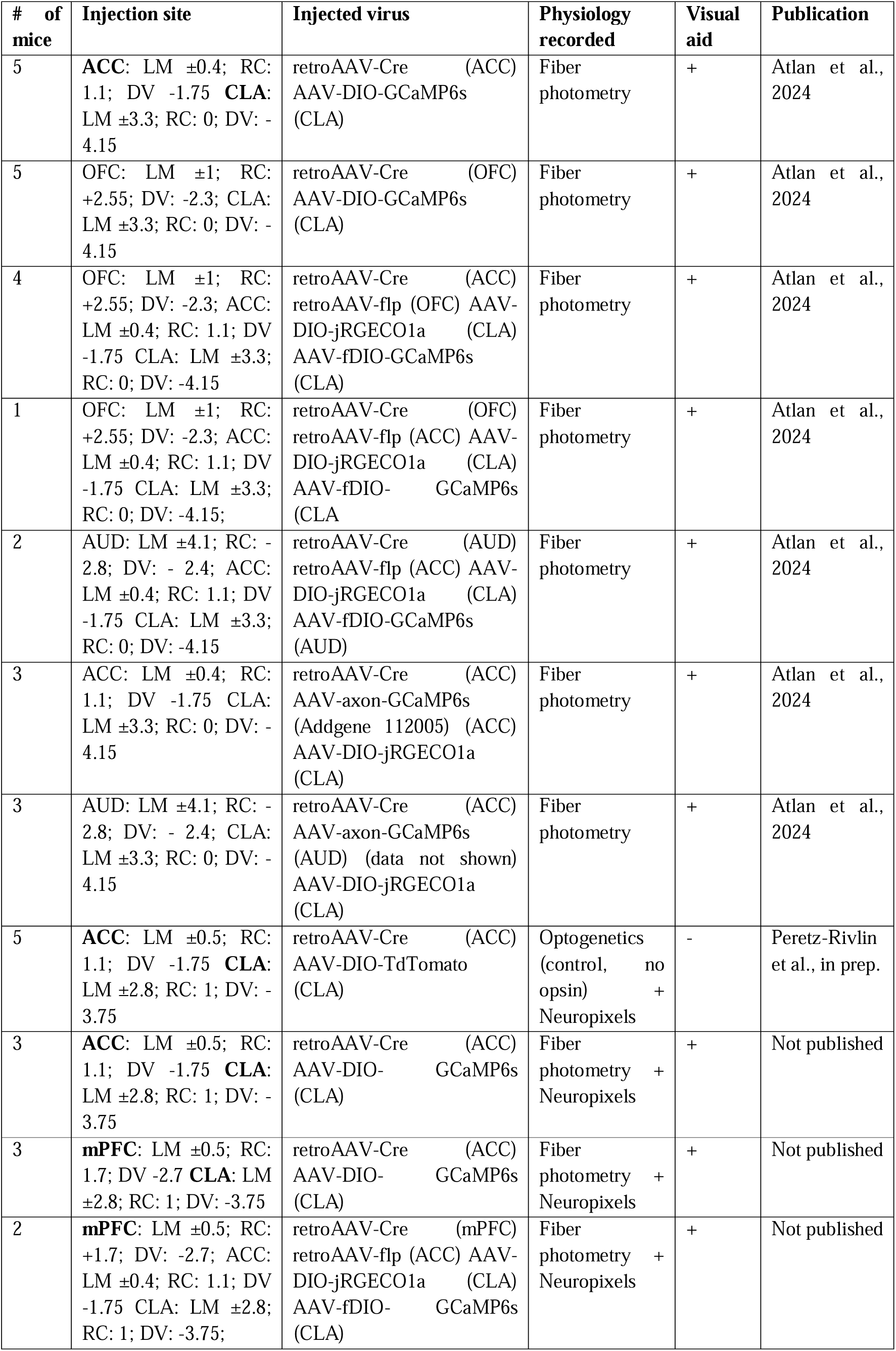

## References

Aguillon-Rodriguez, V., Angelaki, D., Bayer, H., Bonacchi, N., Carandini, M., Cazettes, F., Chapuis, G., Churchland, A. K., Dan, Y., Dewitt, E., Faulkner, M., Forrest, H., Haetzel, L., Häusser, M., Hofer, S. B., Hu, F., Khanal, A., Krasniak, C., Laranjeira, I., … Zador, A. M. (2021). Standardized and reproducible measurement of decision-making in mice. ELife, 10. 10.7554/eLife.63711

Ashwood, Z. C., Roy, N. A., Stone, I. R., Urai, A. E., Churchland, A. K., Pouget, A., & Pillow, J. W. (2022). Mice alternate between discrete strategies during perceptual decision-making. Nature Neuroscience, 25(2), 201–212. 10.1038/s41593-021-01007-z

Atlan, G., Matosevich, N., Peretz-Rivlin, N., Marsh-Yvgi, I., Zelinger, N., Chen, E., Kleinman, T., Bleistein, N., Sheinbach, E., Groysman, M., Nir, Y., & Citri, A. (2024). Claustrum neurons projecting to the anterior cingulate restrict engagement during sleep and behavior. Nature Communications, 15(1). 10.1038/s41467-024-48829-6

Bari, A., & Robbins, T. W. (2013). Inhibition and impulsivity : Behavioral and neural basis of response control. Progress in Neurobiology, 108, 44–79. 10.1016/j.pneurobio.2013.06.005

Berditchevskaia, A., Cazé, R. D., & Schultz, S. R. (2016). Performance in a GO/NOGO perceptual task reflects a balance between impulsive and instrumental components of behaviour. Scientific Reports, 6. 10.1038/srep27389

Bissonette, G. B., & Powell, E. M. (2012). Reversal learning and attentional set-shifting in mice. Neuropharmacology, 62(3), 1168–1174. 10.1016/j.neuropharm.2011.03.011

Bogacz, R., Brown, E., Moehlis, J., Holmes, P., & Cohen, J. D. (2006). The Physics of Optimal Decision Making : A Formal Analysis of Models of Performance in Two-Alternative Forced-Choice Tasks. 113(4), 700–765. 10.1037/0033-295X.113.4.700

Busse, L., Ayaz, A., Dhruv, N. T., Katzner, S., Saleem, A. B., Schölvinck, M. L., Zaharia, A. D., & Carandini, M. (2011). The detection of visual contrast in the behaving mouse. Journal of Neuroscience, 31(31), 11351–11361. 10.1523/JNEUROSCI.6689-10.2011

Carandini, M., & Churchland, A. K. (2013). Probing perceptual decisions in rodents. In Nature Neuroscience (Vol. 16, Issue 7, pp. 824–831). 10.1038/nn.3410

Gilad, A., Gallero-Salas, Y., Groos, D., & Helmchen, F. (2018). Behavioral Strategy Determines Frontal or Posterior Location of Short-Term Memory in Neocortex. Neuron, 99(4), 814–828.e7. 10.1016/j.neuron.2018.07.029

Gouveia, K., & Hurst, J. L. (2017). Optimising reliability of mouse performance in behavioural testing: The major role of non-aversive handling. Scientific Reports, 7. 10.1038/srep44999

Grieco, F., Bernstein, B. J., Biemans, B., Bikovski, L., Burnett, C. J., Cushman, J. D., van Dam, E. A., Fry, S. A., Richmond-Hacham, B., Homberg, J. R., Kas, M. J. H., Kessels, H. W., Koopmans, B., Krashes, M. J., Krishnan, V., Logan, S., Loos, M., McCann, K. E., Parduzi, Q., … Noldus, L. P. J. J. (2021). Measuring Behavior in the Home Cage: Study Design, Applications, Challenges, and Perspectives. In Frontiers in Behavioral Neuroscience (Vol. 15). Frontiers Media S.A. 10.3389/fnbeh.2021.735387

Guo, Z. V., Li, N., Huber, D., Ophir, E., Gutnisky, D., Ting, J. T., Feng, G., & Svoboda, K. (2014). Flow of cortical activity underlying a tactile decision in mice. Neuron, 81(1), 179–194. 10.1016/j.neuron.2013.10.020

Harlow, H. F. (1949). THE FORMATION OF LEARNING SETS. Psychological Review, 56, 51–65.

Murphy, T. H., Michelson, N. J., Boyd, J. D., Fong, T., Bolaños, L. A., Bierbrauer, D., Siu, T., Balbi, M., Bolaños, F., Vanni, M., & Ledue, J. M. (2020). Automated task training and longitudinal monitoring of mouse mesoscale cortical circuits using home cages. ELife, 9, 1–91. 10.7554/eLife.55964

Pan, S. J., & Yang, Q. (2010). A Survey on Transfer Learning. IEEE Transactions on Knowledge and Data Engineering, 22(10), 1345–1359. 10.1109/TKDE.2009.191

Peretz-Rivlin, N., Marsh-Yvgi, I., Fatal, Y., Terem, A., Turm, H., Shaham, Y., & Citri, A. (2024). An automated group-housed oral fentanyl self-administration method in mice. Psychopharmacology. 10.1007/s00213-024-06528-6

Poddar, R., Kawai, R., & Ölveczky, B. P. (2013). A fully automated high-throughput training system for rodents. PLoS ONE, 8(12). 10.1371/journal.pone.0083171

Raposo, D., Sheppard, J. P., Schrater, P. R., & Churchland, A. K. (2012). Multisensory decision-making in rats and humans. Journal of Neuroscience, 32(11), 3726–3735. 10.1523/JNEUROSCI.4998-11.2012

Shenhav, A., Botvinick, M. M., & Cohen, J. D. (2013). The Expected Value of Control: An Integrative Theory of Anterior Cingulate Cortex Function. Neuron, 79(2), 217–240. 10.1016/j.neuron.2013.07.007

Siniscalchi, M. J., Phoumthipphavong, V., Ali, F., Lozano, M., & Kwan, A. C. (2016). Fast and slow transitions in frontal ensemble activity during flexible sensorimotor behavior. Nature Neuroscience, 19(9), 1234–1242. 10.1038/nn.4342

